# A molecularly-defined non-redundant subpopulation of OPCs controls the generation of myelinating oligodendrocytes during postnatal development

**DOI:** 10.1101/2023.07.28.550937

**Authors:** Shayan Moghimyfiroozabad, Maela A. Paul, Lea Bellenger, Fekrije Selimi

## Abstract

Oligodendrocyte precursor cells (OPCs) are a class of glial cells that uniformly tiles the whole central nervous system. They play several key functions across the brain including the generation of oligodendrocytes and the control of myelination. Whether the functional diversity of OPCs is the result of genetically defined subpopulations or of their regulation by external factors has not been definitely established. We discovered that a subpopulation of OPCs found across the brain is defined by the expression of *C1ql1*, a gene previously described for its synaptic function in neurons. This subpopulation starts to appear during the first postnatal week in the mouse brain. Ablation of *C1ql1*-expressing OPCs in the mouse is not compensated by the remaining OPCs, and results in a massive lack of oligodendrocytes and myelination in many brain regions. Therefore, *C1ql1* is a molecular marker of a functionally non-redundant subpopulation of OPCs, which controls the generation of myelinating oligodendrocytes.

## Introduction

Oligodendrocyte precursor cells (OPCs) are a type of glial cells that homogeneously populate the central nervous system (CNS) starting during postnatal development and throughout adulthood^1^. Several functions have been attributed to OPCs^2^. OPCs have been documented to control vascularization and angiogenesis^3^, engulfment of axons and refinement of neural circuits^4,5^, and antigen presenting^6^. However, a major, and their initially described, role is to differentiate into oligodendrocytes (OLs)^7–9^. In rodents, the generation of OLs from OPCs starts at birth and continues during adulthood^10,11^. OPC differentiation peaks during the second postnatal week and then decreases gradually^10^. A portion of OPCs continues to proliferate to keep the density of their population constant throughout the mature brain^1^. OPCs are thus essential for proper development and function of brain circuits. OPCs are characterized by the expression of neural/glial antigen 2 (NG2) and platelet-derived growth factor receptor A (PDGFRa), which immediately disappear upon differentiation into OLs^1,9^.

OPCs are generated in the CNS through sequential waves and in a ventro-dorsal manner^7^. In the forebrain, three waves with different spatial origins and genetic lineages generate OPCs^12^. The first two waves are originally ventral and embryonic. The third wave is generated dorsally after birth^12^. The elimination of each wave of OPCs is rapidly compensated by the other waves without any significant consequences on OPC dynamics or myelination of the developing brain^12^. Even the ablation of all three waves in the forebrain is compensated by OPCs coming from the diencephalon within two weeks^7,13^. These data suggest that, despite their different origins, various waves of OPCs are functionally redundant. However, functional data also suggest heterogeneity in OPCs ^14^. For example, OPCs coming from the first wave in the forebrain migrate towards the optic nerve via a Nrg1-ErbB4-mediated chemo-affinity mechanism, while OPCs from the 2^nd^ wave that occupy the optic nerve postnatally are Nrg1-insensitive^15^. Another example is the modulation of the generation of first wave OPCs via Shh signaling, while the other waves are Shh-independent^16^. Independent of their origin, OPCs have slightly different morphologies based on their location^1,17^. For example, OPCs in the white matter have an elongated soma packed tightly between the myelinated axons, with extended processes in parallel to the direction of the fibers. OPCs in the gray matter have a star-shape morphology with several long and thin branches extending in all directions^1^. OPCs can also have different dynamics in terms of proliferation, migration and differentiation rate^8^. For example, OPCs in the corpus callosum differentiate into oligodendrocytes faster than OPCs in the cortex^8^. OPCs are also heterogeneous based on their transcriptomes, cell surface proteins (e.g. receptors and ion channels) or electrophysiological properties^17–19^. Single cell RNA-seq experiment performed on OPCs coming from mouse forebrain at different stages during development (E13.5, P7, and P21) shows that OPCs originating from several regions of the forebrain become transcriptionally equivalent at P7^20^. Despite this transcriptional equivalence, OPCs in ventral and dorsal forebrain respond differently to DNA damage by promoting senescence or apoptosis, respectively. These two different responses are linked to different molecular mechanisms and developmental origin of OPCs, showing molecular and functional heterogeneity in OPCs^21^. In addition to heterogeneity associated with the origin, OPC heterogeneity can change with age too^19^. Spitzer *et al.* showed that OPCs lack any ion channels at E13.5 in different regions of the brain, while they start to express voltage-gated ion channels and glutamate receptors differentially throughout development in concordance with their functional states^19^. All these studies demonstrate that OPCs are a diverse population and their diversity changes based on the brain region and age.

A fundamental question is whether the functional diversity in OPCs arises from molecularly defined subpopulations with distinct functions or whether it is a result of their interaction with the environment^1,17^. In this study, we show that *C1ql1*, a gene known for its role in neuronal synapse formation^22,23^, is a molecular marker of a subpopulation of OPCs all over the mouse brain starting during the first postnatal week. Using genetic ablation, we show that the elimination of this subpopulation is not compensated by the remaining OPCs and leads to lack of OLs in the dorsal forebrain. Consequently, the myelination of the brain is massively decreased. Thus, for the first time, we identify a molecular signature for a non-redundant subpopulation of OPCs that generates myelinating OLs during the postnatal development of mouse forebrain.

## Results

### A subpopulation of OPCs is characterized by *C1ql1* expression

The *C1ql1^Cre^* knockin mouse model^24^ can be used to determine the history of expression of *C1ql1* during brain development. For this we used the Cre-dependent *R26^Cas9-GFP^* (known as Rosa26-Cas9 knockin)^25^ reporter mouse line allowing simultaneous expression of the CAS9 protein and the green fluorescent protein (GFP) upon Cre-mediated recombination (Figure 1A). In addition to the expected expression in certain neuronal populations^24^, we noticed the presence of GFP-expressing cells throughout the brain at postnatal day 30 (P30) (Figure 1B). These cells did not appear to be neurons, an observation that was confirmed using immunolabeling for NeuN (Figure S1). Indeed, while cerebellar granule cells and inferior olivary neurons were both GFP and NeuN positive as expected (Figure S1), the other GFP-labeled cells were NeuN negative. Because the morphology of these cells was suggestive of glia, we tested markers for the various glial populations. The GFP-positive cells were not labeled using an anti-GFAP antibody showing that they were not astrocytes (Figure S1). Using NG2, a well characterized marker of oligodendrocyte precursor cells (OPCs), we determined that the non-neuronal cells labeled using the *C1ql1^Cre^*knockin mouse line were a subpopulation of OPCs: some, but not all, NG2 labeled cells were also GFP-positive (Figure 1B).

**Figure 1.**
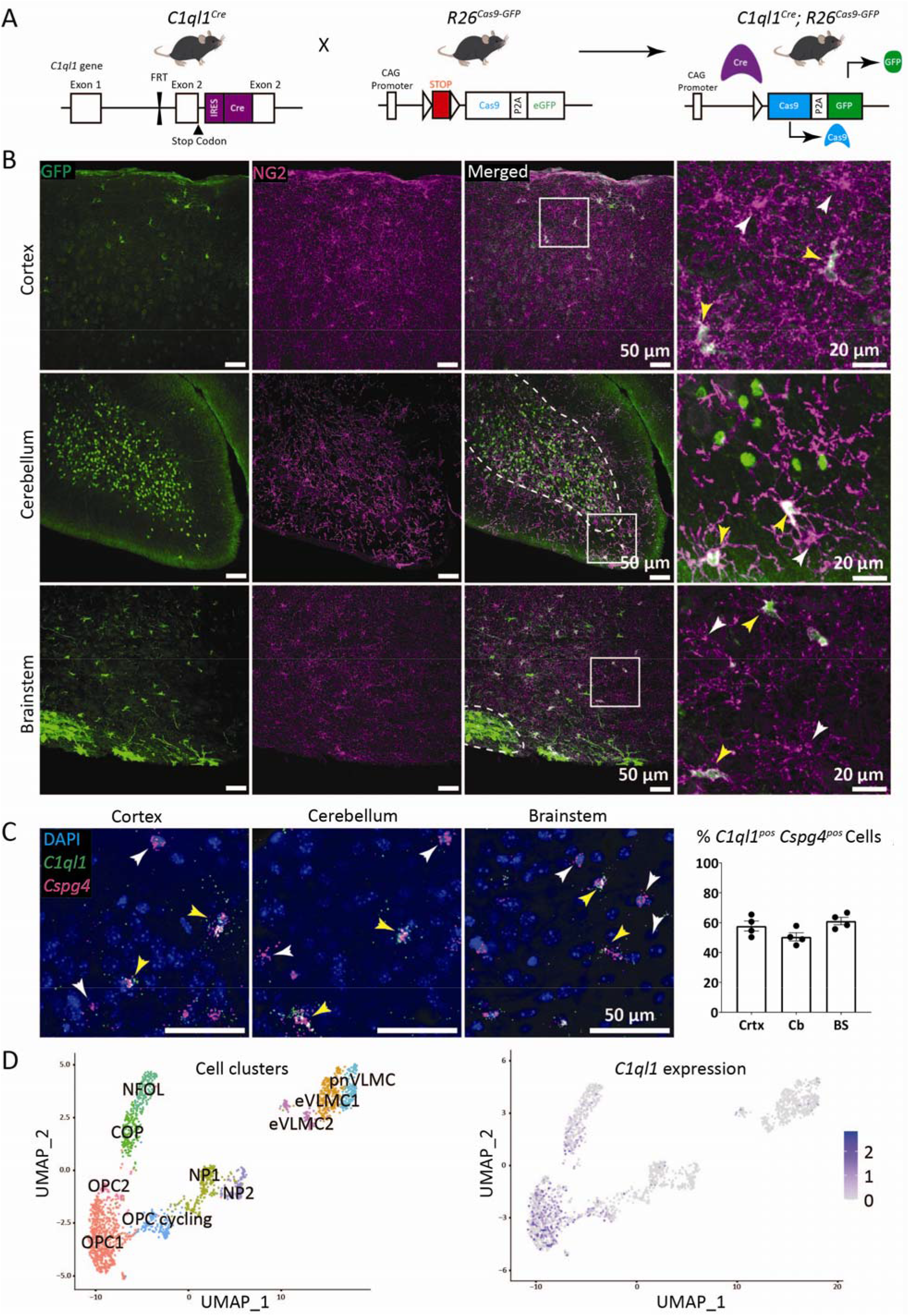
A subpopulation of Oligodendrocyte Precursor Cells is characterized by *C1ql1* expression. A) Diagram of the breeding scheme used to determine the cell types expressing *C1ql1* in the mouse brain. The *C1ql1^Cre^* mouse line was crossed with the Cre-dependent *R26^Cas9-GFP^* reporter mice. B) GFP-expressing cells and Oligodendrocyte Precursor Cells (OPCs) were visualized using co-immunolabeling for GFP (green) and NG2 (magenta) respectively, in sagittal brain sections from *C1ql1^Cre^; R26^Cas9-GFP^* animals at postnatal day 30 (P30). In the cerebellum and the brainstem, GFP is strongly expressed by granule cells and inferior olivary neurons, respectively (internal granular layer and inferior olive delineated by the white dashed line). Scale bars = 50 µm. Higher magnification of the regions delineated by the white squares show the presence of GFP^neg^ (white arrowhead) and GFP^pos^ (yellow arrowhead) NG2 cells in all analyzed regions of the brain. Scale bars = 20 µm. C) Duplex smFISH labeling of *C1ql1* and *Cspg4* (coding for NG2) mRNAs was performed in parasagittal sections of the somatosensory cortex, cerebellum, and brainstem from P30 wild-type animals. Both *C1ql1^neg^*(white arrowhead) and *C1ql1^pos^* (yellow arrowhead) *Cspg4^pos^*cells are detected. Scale bars = 50 µm. Quantification of the percentage of *C1ql1^pos^ Cspg4^pos^* cells was performed in the somatosensory cortex (Crtx), cerebellum (Cb) and brainstem (BS). Data are represented as mean ± SEM. n = 4 animals. D) New analysis of the single cell transcriptomic raw data from Marques *et al.*^20^ reveals the different cell clusters of the oligolineage (*left panel*) and confirms the expression of *C1ql1* (*right panel*) in a subset of OPCs and COP cells. COP: committed oligodendrocyte precursors, eVLMC: embryonic vascular and leptomeningeal cells, NFOL: newly formed oligodendrocyte, NP: neural progenitor, OPC: oligodendrocyte precursor cell, pnVLMC: postnatal vascular and leptomeningeal cells.

This genetic approach indicates that a subpopulation of OPCs, or their progenitors, express *C1ql1*. To determine whether OPCs themselves express *C1ql1*, we performed duplex single molecule fluorescence in situ hybridization (smFISH) for *C1ql1* together with *Cspg4* (coding the marker NG2) on brain sections from P30 mice (Figure 1C). *C1ql1* puncta were observed together with *Cspg4* puncta in a subset of cells throughout the brain. Quantification showed that about 57%, 50%, and 61% of *Cspg4* expressing cells in the cortex, cerebellum, and brainstem respectively, express *C1ql1* (Figure 1C). Single cell transcriptomics of the oligodendrocyte lineage at embryonic (E13.5) and postnatal stages (P7, juvenile and adult) has previously shown that, besides a subpopulation of cycling OPCs, OPCs form a rather homogenous cell population at the transcriptome level at P7^20^. Using the raw expression data obtained in these previous studies^20^, we performed our own single cell transcriptomics analysis to determine the pattern of *C1ql1* expression across the different cell clusters constituting the oligodendrocyte lineage. We used a batch correction to eliminate the effects due to the different experimental batches (E13.5 and P7 vs juvenile and adult cf. Figure S2), followed by uniform manifold approximation and projection (UMAP) reduction and clustering on shared K-nearest neighbor (SNN) graph. This allowed us to identify ten clusters including three OPC subpopulations (Figure 1D). Two of these subpopulations were the same as the ones found by Marques *et al.*^20^ (OPC1 and the cycling OPC subpopulations), while a small third one (OPC2) was defined by the enriched expression of genes associated with the GO terms “mRNA processing” and “mRNA splicing”, suggesting a specific metabolism of RNA. *C1ql1*-expressing cells were found in all three OPC clusters as well as in a subset of committed OPCs (COPs) that are differentiating into oligodendrocytes (Figure 1D). *C1ql1* expression was almost undetected in newly formed oligodendrocytes (NFOL).

Thus, genetic and transcriptomic data altogether demonstrate that a subpopulation of OPCs expresses *C1ql1* in the juvenile brain, and raise the question as to whether this subpopulation of OPCs has a specific role in the brain.

### Ablation of *C1ql1*-expressing OPCs is not compensated during postnatal development

In the adult mouse brain, OPCs maintain their homeostasis by local proliferation^26^. Ablation, differentiation, or apoptosis of OPCs leads to proliferation and migration of the adjacent cells to replace the missing ones^26^. OPCs that are generated in different waves show functional redundancy, as ablation of one wave using *Sox10*-driven expression of diphtheria toxin (DTA) is compensated by the other two waves in the cortex during the first two postnatal weeks^12^. Even ablation of all three waves induces repopulation by the OPCs from other regions of the brain (e.g. diencephalon) within a few weeks, showing the strong capacity of OPCs for sensing OPC disappearance and compensating for it^12,13,26^. We used the previously established DTA-based ablation method to test whether *C1ql1*-expressing OPCs (*C1ql1^pos^*OPCs) are functionally redundant with other OPCs. For this, the *C1ql1^Cre^*mouse line was crossed with the *Sox10^DTA^* mouse line expressing DTA in a Cre dependent manner^12^ (Figure 2A). In the absence of Cre, all the oligolineage cells, OPC, COPs and OLs, express GFP in the *C1ql1^wt^; Sox10^DTA^* mouse line. *C1ql1* is not expressed in progenitors of OPCs (Figure 1D). Thus, the expression of the Cre recombinase using the *C1ql1^Cre^*locus induces DTA expression and death of the Cre-expressing cells in the oligolineage without affecting *C1ql1*-expressing neurons. The *C1ql1^Cre^; Sox10^DTA^* mice did not survive beyond three to four postnatal weeks. All the animals survived until P16, the age chosen for analysis, without any major visually detectable behavior or motor deficit. Morphological analysis of coronal and parasagittal brain sections showed a massive reduction in the number of GFP-expressing cells all over the brain, and in particular in the forebrain, in *C1ql1*-expressing OPC-ablated (*C1ql1^Cre^; Sox10^DTA^*) animals compared to *C1ql1^wt^; Sox10^DTA^*controls (Figure 2B, 2C and 2D). Since the development of oligolineage cells is well-known in the forebrain, particularly in the corpus callosum and the cortex^12^, we selected the central and lateral parts of the corpus callosum (CC-center & CC-lateral), as well as the retrosplenial area of the cortex for detailed analysis (Figure 2C-D). Quantification of the number of *Sox10*-GFP^positive^ (oligolineage) cells shows a 66% decrease in the center (mean ± SEM = 41.2 ± 2.1 vs 13.9 ± 1.2 x 10^-5^ cells/μm^3^ in the control and ablation conditions, respectively) and 85% in the lateral (39.1 ± 3.0 vs 5.7 ± 0.6 x 10^-5^ cells/μm^3^) parts of the corpus callosum, while a 46% decrease is measured in the cortex (10.0 ± 1.5 vs 5.4 ± 0.5 x 10^-5^ cells/μm^3^) (Figure 2C). Since GFP labels the entire oligolineage cells in the *Sox10^DTA^* mouse model, the OPC marker NG2^1,9^ was used to quantify the OPC subpopulation (NG2^Pos^ GFP^Pos^). A 57 and 80% reduction in the density of NG2^+^ GFP^+^ cells was found in the center and lateral parts of the corpus callosum, respectively (25.0 ± 4.1 vs 10.7 ± 1.0 and 24.3 ± 3.2 vs 4.8 ± 0.6 x 10^-5^ cells/μm^3^, respectively). A 32% decrease was also observed in the cortex (mean ± SEM = 7.4 ± 1.6 vs 5.0 ± 0.5 x 10^-5^ cells/μm^3^). In comparison with NG2 staining, immunolabeling for PDGFRa, another marker of OPCs^1,9^, is more concentrated on the somata of OPCs and allows easier quantification of individual OPCs (Figure 2C-D). Our quantifications found a 74% (mean ± SEM = 20.1 ± 1.2 vs 5.3 ± 0.7 x 10^-5^ cells/μm^3^) and 79% (26.5 ± 3.3 vs 5.7 ± 0.4 x 10^-5^ cells/μm^3^) decrease in the density of PDGFRa^+^ GFP^+^ cells in the central and lateral parts of the corpus callosum, respectively, and a 29% reduction in the cortex (mean ± SEM = 7.9 ± 0.8 vs 5.6 ± 0.4 x 10^-5^ cells/μm^3^). Altogether, using the two most common markers of OPCs, NG2 and PDGFRa, we confirm that DTA expression using the *C1ql1* knockin as a driver leads to the significant ablation of OPCs in several regions of the forebrain. This result was completely unexpected as it also shows that remaining OPCs cannot compensate for the loss of *C1ql1*-expressing OPCs during postnatal development, contrary to what is seen when ablating OPCs from different waves for example^12,13^.

**Figure 2.**
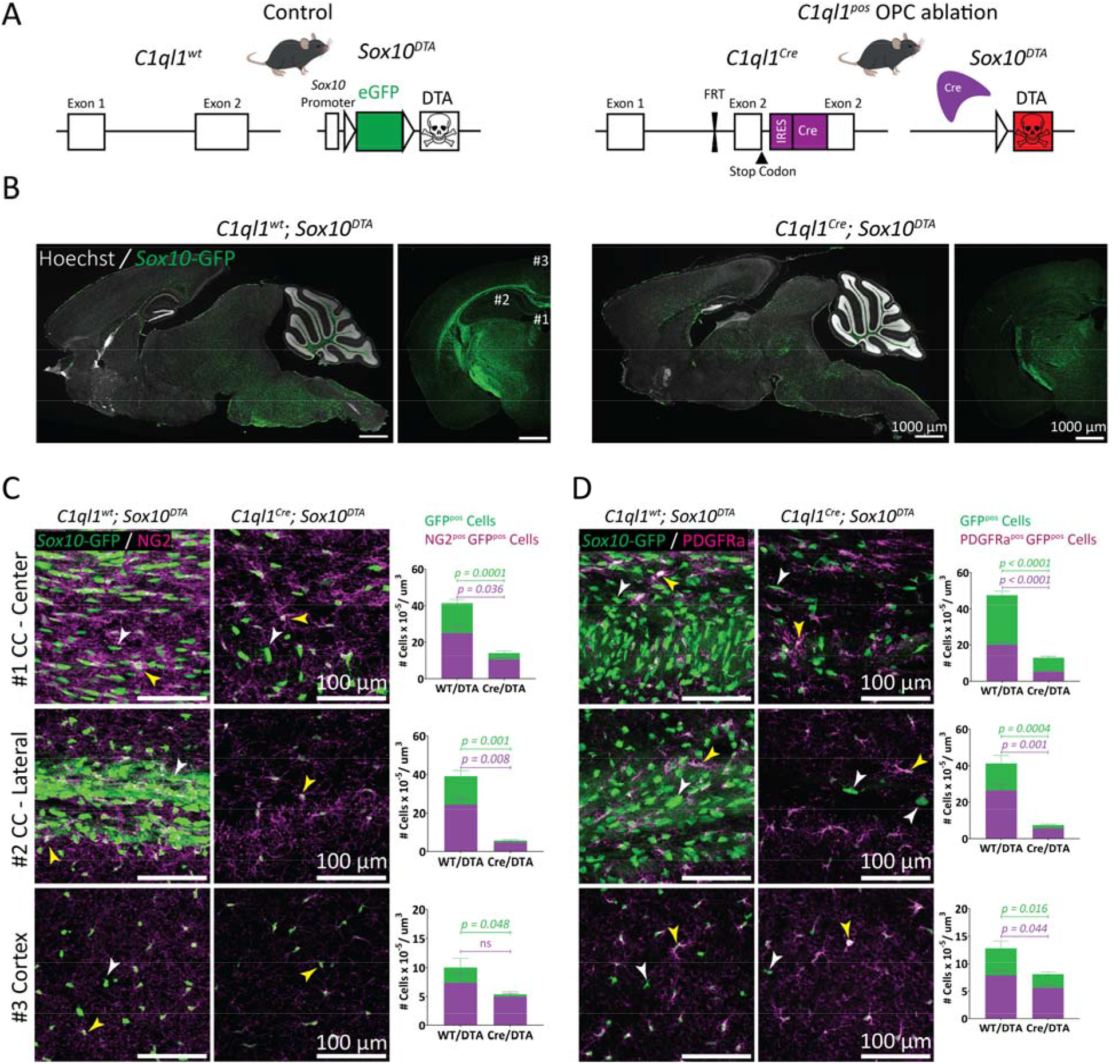
Ablation of *C1ql1*-expressing OPCs is not compensated by remaining OPCs during postnatal development. A) Diagram of the breeding scheme used to ablate *C1ql1*- expressing oligolineage cells in the mouse brain via *Sox10*-driven expression of diphtheria toxin (DTA). The *C1ql1^Cre^* mouse line was crossed with the Cre-dependent *Sox10^DTA^* mice. B) Parasagittal and coronal sections from P16 brains of control (*C1ql1^wt^; Sox10^DTA^*) and *C1ql1*^pos^ OPC-ablated (*C1ql1^Cre^; Sox10^DTA^*) mice. Direct GFP fluorescence and Hoechst staining are shown. Scale bars = 1000 µm. C) OPCs were immunolabeled for NG2 (magenta) in coronal sections from P16 brains of the control and *C1ql1*-ablated conditions. Both NG2^pos^ (yellow arrowhead) and NG2^neg^ (white arrowhead) *Sox10*-GFP^pos^ cells are detected in the center and lateral part of corpus callosum (CC) as well as the cortex, but in lower number in the ablated condition. Scale bars = 100 µm. Quantification of the density of *Sox10*-GFP (green) and NG2^+^- *Sox10*-GFP (magenta) was performed in CC-center, CC-lateral and cortex from the two genotypes. Data are represented as mean ± SEM. *C1ql1^wt^; Sox10^DTA^*: n = 4 animals. *C1ql1^Cre^; Sox10^DTA^*: n = 7 animals. p values are from Unpaired t test with Welch’s corrections. D) OPCs were immunolabeled for PDGFRa (magenta) in coronal sections from P16 brains of the control and *C1ql1^Cre^; Sox10^DTA^* mice. Like for NG2 staining, both PDGFRa^pos^ (yellow arrowhead) and PDGFRa^neg^ (white arrowhead) *Sox10*-GFP^pos^ cells are observed in the two groups in the center and lateral part of corpus callosum (CC) as well as the cortex, but with different ratios. Scale bars = 100 µm. Quantification of the density of *Sox10*-GFP^pos^ (green) and PDGFRa^pos^ *Sox10*- GFP^pos^ (magenta) in the two genotypes confirms the decrease in the number and ratio of the oligolineage cells and OPCs upon ablation of *C1ql1*-expressing *Sox10^pos^*cells. Data are represented as mean ± SEM. *C1ql1^wt^; Sox10^DTA^*: n = 6 animals. *C1ql1^Cre^; Sox10^DTA^*: n = 10 animals. p values are from Unpaired t test with Welch’s corrections. ns = not significant.

### *C1ql1* expression in OPCs starts from the first postnatal week and leads to different dynamics in the dorsal forebrain

The magnitude of OPC loss at P16 in the *C1ql1^Cre^; Sox10^DTA^* mice is bigger in the corpus callosum than in the cortex. In the cortex, 30% of OPCs are missing, a percentage inferior to the one of OPCs expressing *C1ql1* by P30 (about 60% as found by smFISH). We wondered whether these regional differences and in particular the smaller loss in the cortex were due to differences in the timing of expression of *C1ql1* in OPCs during postnatal development. Analysis of previously published data of single cell transcriptomics of oligolineage cells^20^ shows a negligible expression of *C1ql1* at E13.5, while a considerable number of OPCs express *C1ql1* already by P7 and in juvenile animals (Figure 3A) suggesting that *C1ql1* expression is induced during the first postnatal week in a subpopulation of OPCs in the forebrain. To quantify *C1ql1* expression during the first two postnatal weeks, we performed duplex smFISH for *C1ql1* and *Cspg4* on brain sections from wildtype mouse at P7 and P15 (Figure 3A). Similar to our observations at P30 (Figure 1C), *C1ql1* labeling was detected in a subgroup of *Cspg4^pos^* cells (Figure 3A). Quantification demonstrated that 72 and 68% of *Cspg4^pos^*cells in the cortex express *C1ql1* at P7 and P15, respectively, a percentage higher by about 10% than the percentage quantified at P30 (Figure 1C). *C1ql1* expression is thus acquired in a subpopulation of OPCs as early as the first week of postnatal development.

**Figure 3.**
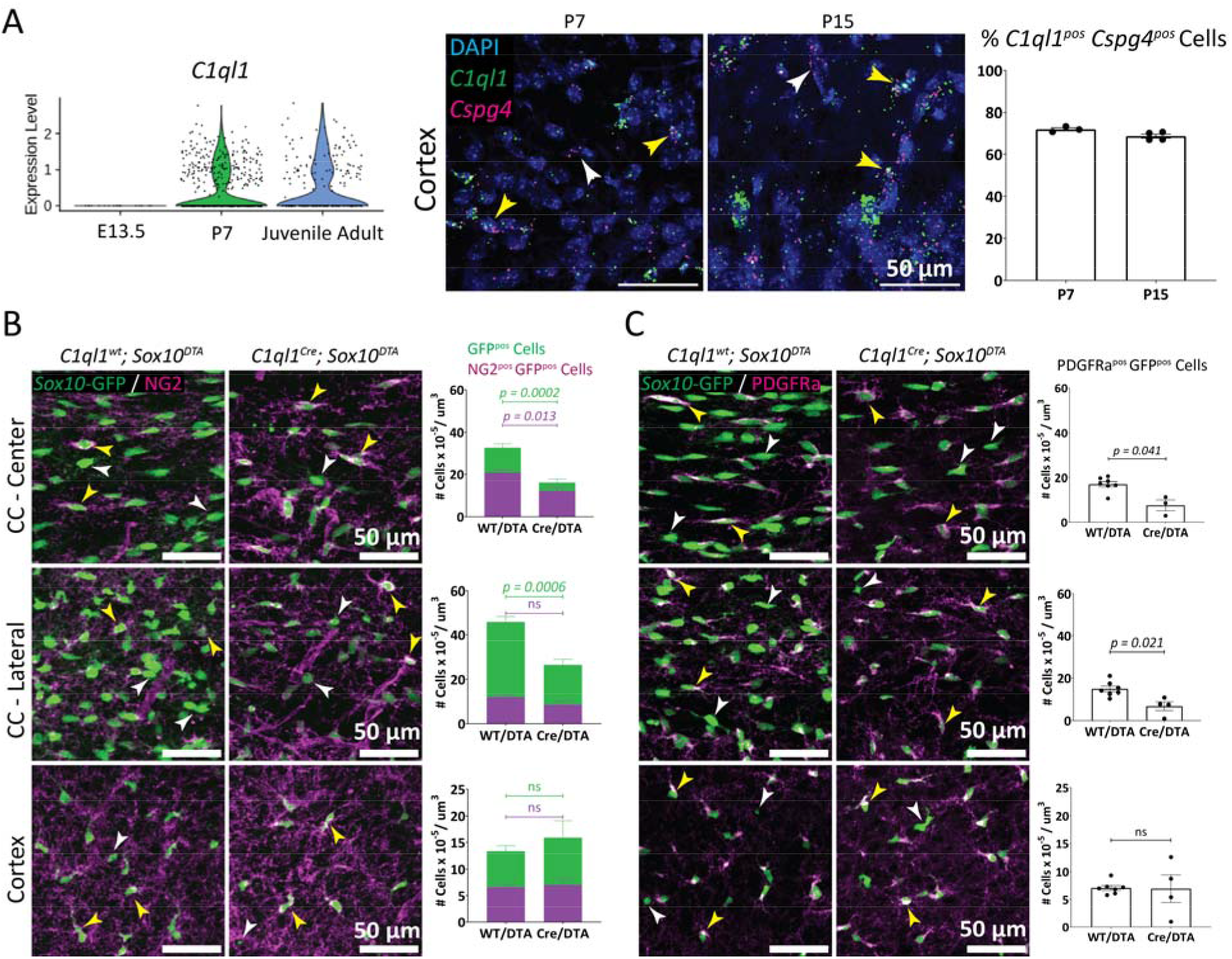
Loss of OPCs starts as early as the first postnatal week in some brain regions after genetic ablation of *C1QL1*-expressing OPCs. A) Violin plot of single cell RNA seq data (from Marques *et al.*^20^) during embryonic and postnatal development shows expression of *C1ql1* as early as P7, but not at E13.5. Duplex smFISH labeling of *C1ql1* and *Cspg4* mRNAs was performed in coronal sections from the cortex of P7 and P15 wild-type mice. Both *C1ql1^neg^* (white arrowhead) and *C1ql1^pos^* (yellow arrowhead) *Cspg4^pos^* cells are observed. Scale bars = 50 µm. Quantification of the percentage of *C1ql1*-expressing *Cspg4^pos^* cells was performed in cortical sections from P7 and P15 animals. Data are represented as mean ± SEM. P7: n = 3 animals. P15: n = 4 animals. Scale bars = 50 µm. B) OPCs were immunostained for NG2 (magenta) in coronal sections from P7 control or *C1ql1^Cre^; Sox10^DTA^*animals. NG2^pos^ (yellow arrowhead) and NG2^neg^ (white arrowhead) *Sox10*-GFP^pos^ cells are identified in the CC-center, CC-lateral and the cortex. Scale bars = 50 µm. Quantification of *Sox10*-GFP^pos^ (green) and NG2^pos^-*Sox10*-GFP^pos^ (magenta) cells in both genotypes shows a decrease in the density of oligolineage cells in CC-center and lateral, but not in the cortex. Data are represented as mean ± SEM. *C1ql1^wt^; Sox10^DTA^*: n = 7 animals. *C1ql1^Cre^; Sox10^DTA^*: n = 3 - 4 animals. Unpaired t test with Welch’s corrections. C) PDGFRa immunolabeling of OPCs (magenta) confirms the decrease in OPC density in both regions of the corpus callosum at P7 after *C1ql1*-expressing OPC ablation. PDGFRa^pos^ (yellow arrowhead) and PDGFRa^neg^ (white arrowhead) *Sox10*-GFP^pos^ cells are observed in both groups. Scale bars = 50 µm. Data are represented as mean ± SEM. *C1ql1^wt^; Sox10^DTA^*: n = 7 animals. *C1ql1^Cre^; Sox10^DTA^*: n = 3 - 4 animals. Unpaired t test with Welch’s corrections. ns = not significant.

In the corpus callosum, the percentage of OPCs expressing *C1ql1* is in the range of the OPC loss observed in *C1ql1^Cre^; Sox10^DTA^*mice at P16. However, in the cortex the OPC loss (30%) is smaller than the percentage of *C1ql1^pos^* OPCs at P7 (60-70%). Quantification showed that in the corpus callosum ablation of *C1ql1^pos^* OPCs has already an effect on the number of oligolineage cells by P7 (Figure 3B-C): The number of GFP cells is already decreased by 40- 50% (Center: mean ± SEM = 32.7 ± 1.9 in the control vs 16.3 ± 1.5 x 10^-5^ cells/μm^3^; Lateral: mean ± SEM = 45.9 ± 2.5 vs 26.6 ± 2.5 x 10^-5^ cells/μm^3^). No change was observed in the cortex at this stage (mean ± SEM = 13.3 ± 1.0 vs 15.9 ± 3.2 x 10^-5^ cells/μm^3^) (Figure 3B). Using NG2 to distinguish OPCs from other oligolineage cells, we observed that the number of NG2^pos^ *Sox10*- GFP^pos^ cells is decreased by 38% (mean ± SEM = 20.1 ± 1.3 vs 12.4 ± 1.7 x 10^-5^ cells/μm^3^) in the center of the corpus callosum, and by 28% in the lateral part of corpus callosum (mean ± SEM = 12.1 ± 0.8 vs 8.7 ± 1.3 x 10^-5^ cells/μm^3^, *p* = 0.0828). No significant changes were detected in the cortex at this stage (mean ± SEM = 6.6 ± 0.5 vs 7.0 ± 1.1 x 10^-5^ cells/μm^3^). Similarly, using PDGFRa as a marker for OPCs, the density of PDGFRa^pos^ *Sox10*-GFP^pos^ cells was found to be decreased by 56% (mean ± SEM = 17.1 ± 1.2 vs 7.6 ± 2.5 x 10^-5^ cells/μm^3^) and 54% (mean ± SEM = 14.9 ± 1.3 vs 6.9 ± 2.1 x 10^-5^ cells/μm^3^) in the center and lateral corpus callosum respectively, while no change was detected in the cortex (mean ± SEM = 7.1 ± 0.4 vs 6.9 ± 2.5 x 10^-5^ cells/μm^3^). These results show that the genetic ablation of *C1ql1*-expressing OPCs has already started during the first postnatal week and is not compensated in the center and lateral part of corpus callosum. In the cortex, OPC loss is not yet observed at the end of the first postnatal week despite the fact that *C1ql1* expression is already detected in OPCs and that a 30% loss is measured one week later (Figure 2C-D). This suggests that in the cortex the loss of OPCs due to ablation is compensated during the first postnatal week by the migration of newly formed OPCs, which is not yet complete at this age^12^. All the OPCs from the 3 different waves are settled in the dorsal forebrain by P10 and their proportions do not change afterwards^12^. Thus, starting at that age, the progressive loss of OPCs in *C1ql1^Cre^; Sox10^DTA^* mice is not compensated by the remaining OPCs during postnatal development.

### Ablation of *C1ql1*-expressing OPCs leads to lack of oligodendrocytes and myelination

*C1ql1*-expressing cells are present in all OPC clusters defined by single cell transcriptomics (Figure 1D). This implies that instead of defining a genetically distinct subgroup, *C1ql1* is a marker of a specific transitional state and/or a specific function within the OPC subpopulation. We performed overrepresentation analysis of the Gene Ontology (GO) terms associated with *C1ql1^pos^* OPCs compared to other OPCs in single cell transcriptomic data (our reanalysis of data from^20^, Figure 1D, Figure 4A and Figure S3A). “Gliogenesis” and “glial cell differentiation” were the two most significantly overrepresented terms (gene ratio = 0.19 and 0.15 respectively) in *C1ql1^pos^* OPCs (Figure 4A and Figure S3A), suggesting that *C1ql1^pos^* OPCs might already be committed to differentiation. Indeed, *C1ql1* expression is also detected in some committed oligodendrocyte precursors (COPs) but disappears in newly formed oligodendrocytes (NFOLs) (Figure 1D). Other markers have been previously found to label a specific transition stage of OPC differentiation in OLs^27–29^. The long noncoding RNA *Pcdh17it* labels newly born immature OLs, as confirmed by its detection mostly in cells from this cluster in our analysis (Figure 1D, Figure S3B). *Gpr17* and *Bcas1* are markers of COPs^28,29^ (Figure 1D and S3B). *C1ql1^pos^* cells do not overlap with cells expressing *Gpr17*, *Bcas1* and *Pcdh17it* markers to a significant extent while many *C1ql1^pos^* cells coexpress *Cspg4*, which labels OPCs (Figure S3B). Thus, *C1ql1* expression is not another marker of COPs and NFOLs, but rather marks a specific state in a subpopulation of OPCs that is ready for differentiation into oligodendrocytes. Indeed, quantification of the number of GFP^pos^ cells after ablation of *C1ql1*-expressing OPCs showed a massive decrease not only in NG2^pos^ GFP^pos^ cells, but also in NG2^neg^ GFP^pos^ cells (Figure 2 and 3). Because GFP expression is driven by the *Sox10* promoter, this suggested that DTA ablation in *C1ql1^pos^* OPCs results in the reduction not only in OPC numbers but also in COPs and oligodendrocytes. Indeed, by P16 NG2^neg^ GFP^pos^ cells are completely absent in the cortex and corpus callosum. Using immunolabeling of P16 forebrain coronal slices for CC1, a marker of oligodendrocytes, we confirmed the massive reduction in OL numbers after ablation of *C1ql1*- expressing OPCs, with almost complete absence of CC1^pos^ cells in the cortex (Figure 4B). Quantification demonstrated a 77%, 98%, and 83% reduction in the number of CC1^pos^ GFP^pos^ cells in the center (mean ± SEM = 18.9 ± 2.3 vs 4.2 ± 1.9 x 10^-5^ cells/μm^3^) and lateral (mean ± SEM = 15.2 ± 1.7 vs 0.2 ± 0.0 x 10^-5^ cells/μm^3^) corpus callosum, and in the cortex (mean ± SEM = 1.2 ± 0.3 vs 0.2 ± 0.1 x 10^-5^ cells/μm^3^), respectively. Thus *C1ql1*-expressing OPCs are essential for the generation and/or survival of OLs.

**Figure 4.**
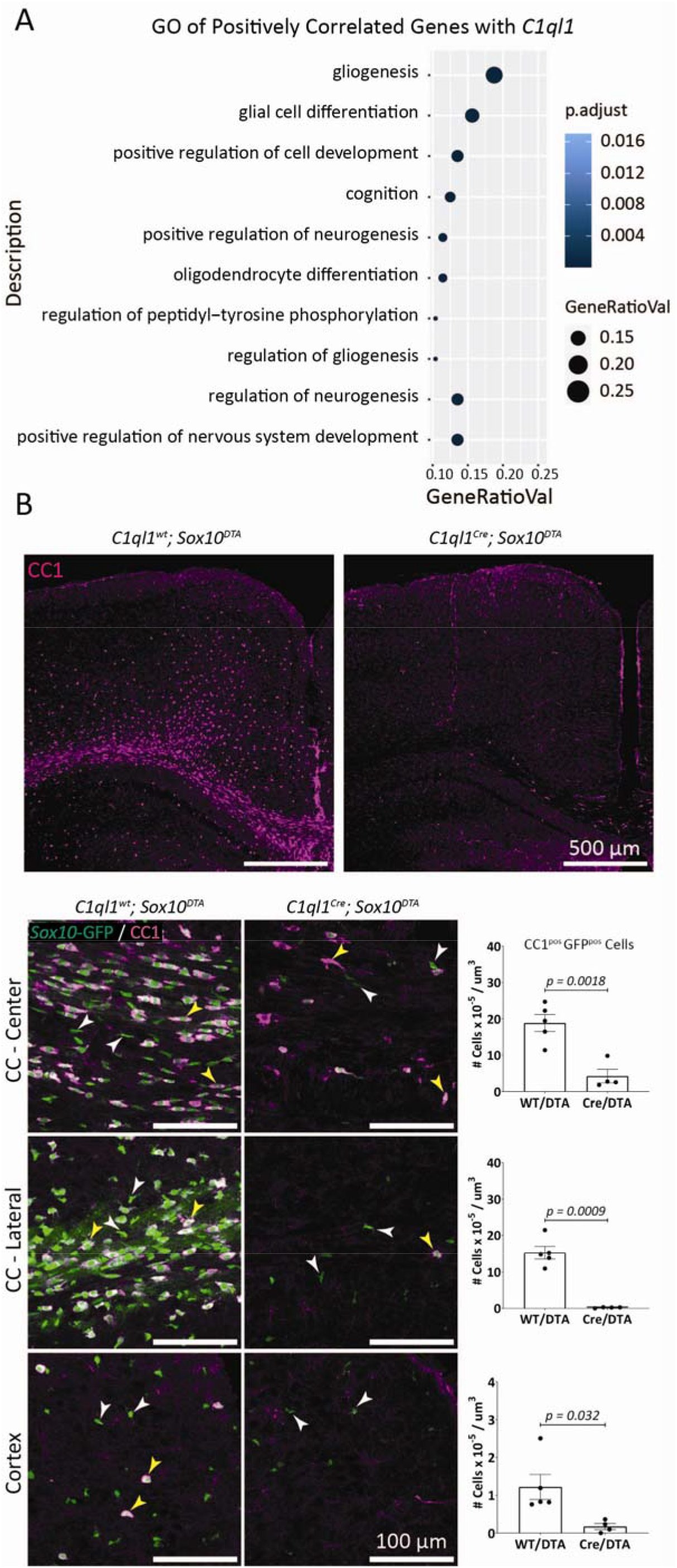
Lack of oligodendrocyte after DTA-mediated ablation of *C1ql1*-expressing OPCs. A) Gene ontology analysis shows the biological functions of the genes enriched in *C1ql1*- expressing OPCs compared to other OPCs (Reanalysis of raw data from Marques *et al.*^20^). B) Low magnification of coronal cortical sections from *C1ql1^wt^; Sox10^DTA^* and *C1ql1^Cre^; Sox10^DTA^* P16 mice immunolabeled with CC1, a marker of oligodendrocytes. Scale bars = 500 µm. Higher magnification of the coronal sections in the center and lateral parts of CC and cortex with CC1 immunolabeling (magenta) shows the dramatic decrease or absence of OLs in *C1ql1^Cre^; Sox10^DTA^* at P16. CC1^pos^ (yellow arrowhead) and CC1^neg^ (white arrowhead) *Sox10*-GFP^pos^ cells are shown. Scale bars = 100 µm. Quantification of the density of *Sox10*-GFP^pos^ (green) and CC1^pos^ *Sox10*-GFP^pos^ (magenta) in the three analyzed regions shows the heterogeneous decrease in OL generation in the *C1ql1*^pos^ OPC-ablated condition. Data are represented as mean ± SEM. *C1ql1^wt^; Sox10^DTA^*: n = 5 animals. *C1ql1^Cre^; Sox10^DTA^*: n = 4 animals. Unpaired t test with Welch’s corrections. ns = not significant.

OL generation starts around birth (P0) in mice and continues throughout life^10^. Myelination starts from the end of the first postnatal week and increases rapidly to reach its maximum level by the end of the second week^10^. This intense period of myelination coincides with the timing of the defects in generation of OLs observed after ablation of *C1ql1*-expressing OPCs. Immunolabeling for myelin binding protein (MBP) revealed a dramatic decrease in MBP protein in different regions of the forebrain at P16 (Figure 5A). Quantification was performed to assess the thickness and intensity of MBP labeling in the corpus callosum and the cortex (Figure 5B). In the center of the corpus callosum, a slight but not statistically significant, decrease was observed in the thickness of the MBP labeled region (mean ± SEM = 320.9 ± 16.8 vs 293.6 ± 12.8 μm), while MBP labeling was almost completely absent in the lateral corpus callosum (mean ± SEM = 144.6 ± 16.8 vs 23.6 ± 4.9 μm) and in the cortex (mean ± SEM = 305.5 ± 52.2 vs 89.6 ± 7.6) (Figure 5B). To rule out an indirect effect of eliminating *C1ql1*-expressing OPCs on axonogenesis, we assessed the integrity of axons and neuronal fibers using immunolabeling for the neurofilament protein (NF)^9,30^ (Figure 5C). No difference was observed in the thickness of fiber tracts in the center of the corpus callosum (mean ± SEM = 326.5 ± 12.3 vs 292.7 ± 17.1 μm) and the intensity of NF labeling in the cortex (mean ± SEM = 742.4 ± 193.1 vs 957.7 ± 208.5). A 19% reduction in the thickness of fiber tracts in the lateral part of the corpus callosum was detected (mean ± SEM = 129.1 ± 7.8 vs 104.9 ± 2.8 μm), and is most likely due to increased compaction of axons rather than to deficits in axonogenesis given the nearly full absence of MBP in this area. In the center of the corpus callosum the effect of ablation of *C1ql1*- expressing OPCs is not obvious as the thickness of MBP and NF labelling remains the same. Some OLs that do not express *Sox10* were observed in the center of corpus callosum (Figure 3B) and it has been reported that 4% of oligodendroglia in the brain are *Sox10*-negative^31^. Since these cells are not eliminated using the *Sox10*-DTA line, they probably compensate for the *Sox10^pos^* lineage in the center of corpus callosum. Overall, our results show a major lack of myelination at P16 in various parts of the forebrain as a consequence of ablation of *C1ql1*- expressing OPCs and lack of oligodendrocytes.

**Figure 5.**
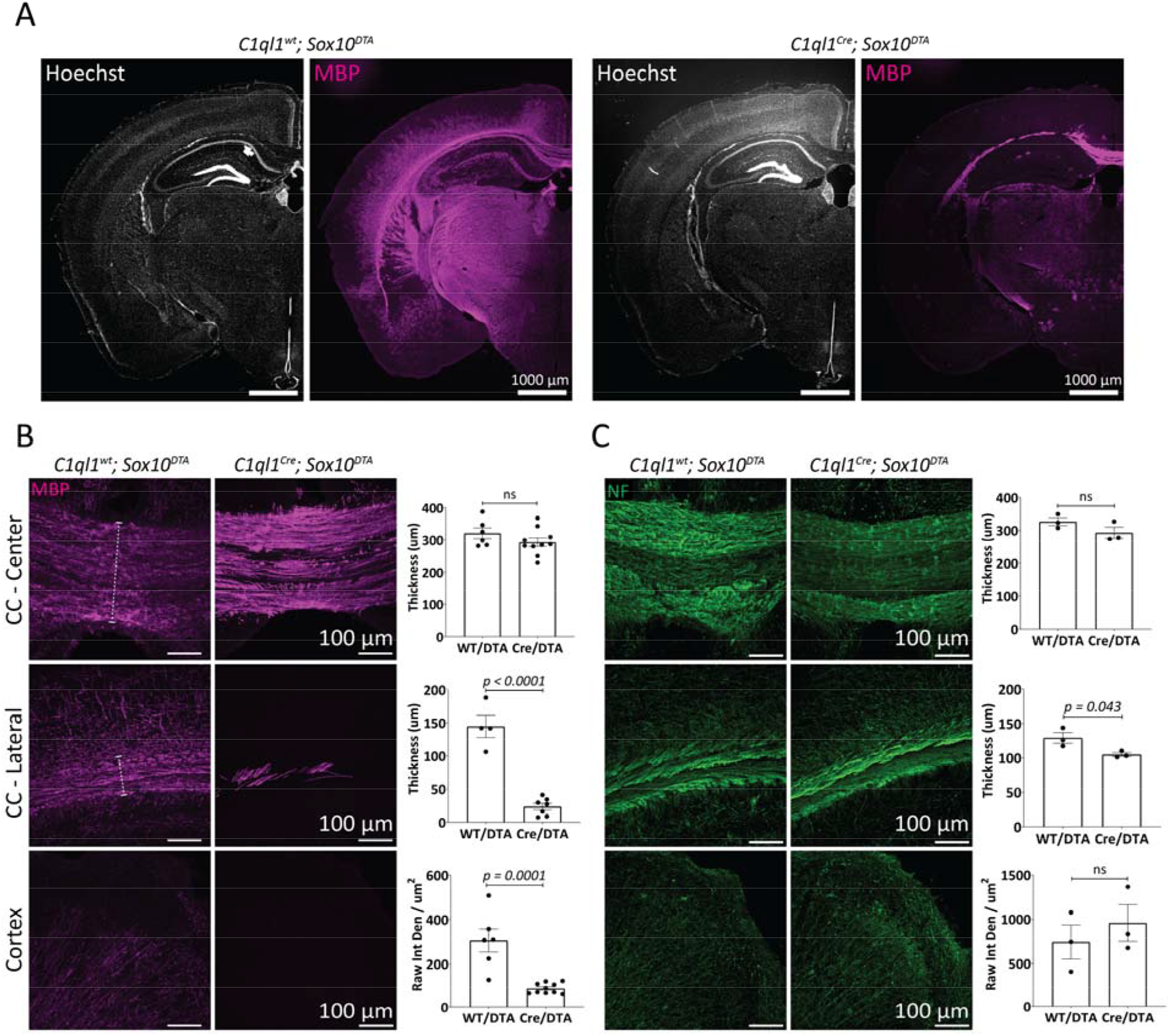
Lack of myelination in the forebrain after genetic ablation of *C1ql1*-expressing OPCs. A) Immunolabeling of coronal brain sections at P16 for myelin binding protein (MBP, magenta) illustrates the almost total absence of myelination in the forebrain upon genetic ablation of *C1ql1*-expressing OPCs, without gross morphological changes. Scale bars = 1000 µm. B) Myelinated fibers (immunolabeled for MBP, magenta) were visualized in coronal sections of the CC-center, lateral, and cortex of the two conditions at P16. The thickness of the myelinated regions in CC, delineated by the white dashed lines, was measured. Scale bars = 100 µm. Quantification of the thickness of myelinated regions in the corpus callosum and the intensity of MBP in the cortex shows an almost complete absence of MBP in CC-lateral and cortex. Data are represented as mean ± SEM. *C1ql1^wt^; Sox10^DTA^*: n = 4 animals. *C1ql1^Cre^; Sox10^DTA^*: n = 7 - 10 animals. Unpaired t test with Welch’s corrections. Raw Int Den: Raw integrated density. C) Neuronal fibers were immunolabeled for neurofilament (NF, green) in coronal P16 sections of the three regions of interest (CC- center / lateral, and cortex), to compare the effect of *C1ql1*-expressing OPC ablation on axogenesis. No deficit is observed when comparing both conditions. Scale bars = 100 µm. Quantification of the thickness of the fiber bundles in the corpus callosum and the intensity of NF in the cortex shows a small decrease in the thickness of the CC-lateral, probably explained by the lack of myelination. Data are represented as mean ± SEM. *C1ql1^wt^; Sox10^DTA^*: n = 3 animals. *C1ql1^Cre^; Sox10^DTA^*: n = 3 animals. Unpaired t test. ns = not significant.

## Discussion

OPCs have various functions^2,14^. Whether this is related to their molecular heterogeneity or is simply the result of interaction with their local environment is unresolved. In this study, we identified a subpopulation of OPCs in the mouse brain defined by the expression of the synaptic molecule *C1ql1* starting during the first postnatal week. Ablation of *C1ql1*-expressing OPCs is not compensated by remaining OPCs in the dorsal forebrain during postnatal development and leads to OL and myelination deficiency.

Single cell RNA sequencing of cells from the oligodendroglia lineage during embryonic and postnatal development previously showed that OPCs with different origins converge transcriptionally at P7 to form a homogeneous population^20^. However, the idea that OPCs are equivalent has been challenged recently^18,21^. One study using single cell transcriptomics on adult mouse OPCs revealed two separate subpopulations of OPCs with distinct molecular markers and enriched in GO terms associated with different biological processes^18^. Another study showed that OPCs from the dorsal forebrain are more vulnerable than the ones from the ventral part to oxidative stress and DNA damage induced by removal of Citron-kinase^21^. Here we present new evidence for molecular and functional diversity of OPCs. The expression of *C1ql1* appears postnatally in a subset of OPCs that reaches their maximum number at P7 and then decreases gradually until P30 in the cortex. Several transcriptomic data show the expression of *C1ql1* is maintained in the adult^18,32^, which suggests the importance of *C1ql1*-expressing OPCs in myelination/re-myelination even after the first postnatal month. In the zebrafish spinal cord, OPCs have been classified into two subpopulations based on several characteristics, including their location, function, calcium activity and dynamics^17^. *C1ql1*-expressing OPCs are found in various regions of the brain and in both white (such as brainstem) and gray matters (e.g. cerebral cortex) in the mouse brain, and *C1ql1* expression is not restricted to a particular cluster of OPCs. Furthermore, there is no significant overlap with markers of COPS and newly formed OLs (NFOLs). Altogether, these data indicate that *C1ql1* marks a transitional state preceding OPC commitment to differentiate into OLs. This state could be induced by interaction of OPCs with the microenvironment or other cells in the tissue. The interaction of OPCs with different types of cells in the brain, such as microglia^2,33^ or endothelial cells^34^, have been shown to be necessary not only for the function of those cells, but also for proper development of OPCs themselves^2,33,34^.

C1QL1 is a synaptic molecule involved in excitatory synapse formation and maintenance between neurons^22,23,35^. Its receptor is the postsynaptic receptor brain angiogenesis inhibitor 3 (BAI3), not only found in neurons^22,23,35^, but also expressed by some glial cells^36^, bringing the possibility that OPCs interact with their local environment, both neurons and other glial cells, via C1QL1-BAI3 signaling. Since OPCs receive synaptic inputs from excitatory neurons^37,38^, it is possible that the C1QL1/BAI3 complex participates in the formation of neuron-OPC connectivity. Additionally, C1QL1 could have a stimulatory effect on vascularization via binding to its receptor BAI3^39^, suggesting a potential role for *C1ql1*-expressing OPCs in angiogenesis. C1QL1 has been proposed to have a proliferative and tumor-promoting function in two different cancers, including one inside the brain^40,41^, and contributes to the regulation of the ovarian follicle reserve^42^. Ablation of *C1ql1*-expressing OPCs leads to the lack of OLs. Thus, an additional role for C1QL1 could be the control of proliferation and/or differentiation of OPCs, especially when an equilibrium between apoptosis and proliferation of oligolineage cells is essential^1,43^.

Demyelination is a pathological condition where the myelin around the axons is lost, due to the damage to the myelin sheath or loss of myelinating OLs. Dysmyelination is another condition where the myelin is not formed properly^44^. Oligolineage cells are at the center of understanding the etiology and treatment of both conditions. For example, in multiple sclerosis (MS), a type of demyelination condition, OPCs have been traditionally studied for the development of remyelination strategies. However, a recent study showed that OPCs might be directly involved in the etiology of the disease: a switch occurs leading to immune like OPCs that are not found in normal brains^45^, and this switch might be directly involved in the etiology of the disease. This switch impairs the ability of OPCs for remyelination^6^. Since no significant change is reported at the level of expression of *C1ql1* in oligolineage cells and the number of *C1ql1*- expressing OPCs upon demyelination in a mouse model of MS using single cell transcriptomics^45^, it seems that this subpopulation is not strongly affected in MS. Due to its importance in myelination during postnatal development, *C1ql1*-expressing OPCs could be a potential target for therapeutic approaches for demyelination/desmyelination conditions.

## Methods and Materials

### Animals

All mice were kept in the authorized animal facilities of CIRB, College de France, under a 12-hour light: 12-hour dark cycle with water and food supplied ad libitum. All animal protocols and animal facilities were approved by the Comité Régional d’Ethique en Expérimentation Animale (#2001) and the veterinary services (D-75-05-12). The *C1ql1^Cre^* mouse model was generated and characterized previously^24^. The Cre-dependent reporter mouse line *R26^Cas9-GFP^*(B6J.129(B6N)-Gt (ROSA)26Sortm1(CAG-cas9*,-EGFP)Fezh/J, strain #026175)^25^ was received from The Jackson Laboratory. All the lines are maintained on the C57BL/6J background. C57Bl6/J and OF1 mice (Charles River Laboratories, Wilmington, USA) were used for the smFISH. *Sox10^DTA^* mice (Sox10-lox-GFP-STOP-lox-DTA, MGI:4999728)^12^ were a kind gift from Pr. William D. Richardson (University College London, UK). They were maintained on a mixed CBA/CaCrl and C57BL/6J background in the original animal facility and were crossed with *C1ql1^Cre^* line upon arrival to CIRB, College de France. In all experiments, littermate mice from both sexes were used as controls for the analysis.

### Single molecule Fluorescent In situ Hybridization (smFISH)

Mice at P7, P15, and P30 were perfused intracardially with 4% paraformaldehyde (PFA) phosphate-buffered saline (PBS) solution. Brains were post-fixed with the same solution at 4°C for 24 hours, transferred to 20% and 30% sucrose/PBS at 4°C sequentially for 24 hours. 30 µm-thick sections (P7 & 15: coronally, P30: parasagittally) were obtained using a freezing microtome and stored at -20°C in cryo-preservative solution until use. The smFISH labelling was performed using RNAscope Multiplex Fluorescent Assay kit (Advanced Cell Diagnostics, Newark, USA, cat#323100) according to manufacturer’s instructions. Duplex in situ hybridization was performed with the *Cspg4* (ACD, cat#404131) and *C1ql1* RNAscope probes (ACD, cat#465081). The nuclei were labelled with DAPI. Sections were mounted with Prolong Gold Antifade reagent (Invitrogen, cat#P36930). Images were taken using a Zeiss spinning-disk confocal CSU-W1 microscope with 40x oil objective. A home-made plugin in ImageJ/Fiji was used to detect and quantify the number of individual RNA puncta and their total surface inside each segmented nucleus. The nuclei with a total surface of the RNA puncta of > 1.5 μm^2^ were selected positive for that probe.

### Immunofluorescence

Mice at P7, P16 - 18, and P30 were fixed using intracardiac perfusion of 4% PFA in PBS solution. Brains were extracted, post-fixed with the same solution at 4°C for 2 - 4 hours, and transferred to 30% sucrose/PBS at 4°C for cryoprotection. 60 µm-thick sections were obtained using a freezing microtome, and kept in 0.02% NaN_3_ in PBS solution at 4°C until use. For CC1 staining (Figure 3B-C), 2% PFA in PBS was used for perfusion and post-fixation.

To perform immunolabeling, slices were incubated in blocking buffer (4% donkey serum and 1% Triton X-100 in PBS solution) for 1 hour at room temperature, followed by incubation with primary antibodies in 1% donkey serum and 1% Triton X-100 in PBS, overnight at 4°C with agitation. Primary antibodies were: GFP (1:1000), chicken, ab13970, Sigma; NG2 (1:1000), rabbit, ab5320, Chemicon; APC/CC1 (1:200), mouse, OP80, Calbiochem; PDGFRa (1:500), rat, 14-1401-81, eBioscience; MBP (1:2000), rat, abcam, ab7349; NF (1:1000), chicken, abcam, ab4680; GFAP (1:500), mouse, G3893, Sigma; NeuN (1:500), rabbit, ab177487, abcam. The slices were washed 3 times in 1% Triton X-100 in PBS for 10 minutes, followed by incubation with secondary antibodies (Alexa Fluor 488-, and 568- and 647-labeled donkey/goat anti-mouse, rat, rabbit, or chicken IgGs (H+L); 1:1000, Invitrogen or Life Technologies) in 1% Triton X-100 in PBS for 2 hours at room temperature. Then, sections were washed 3 times in 1% Triton X-100 in PBS for 10 minutes, and incubated for another 10 min at room temperature with Hoechst 33342 (0,2 mg/mL, Sigma, Gothenburg, Sweden, cat#H6024) and 0.4% Triton X-100 in PBS. Sections were mounted using ProLong Gold Antifade Reagent (Invitrogen, cat#P36930).

### Image acquisition and analysis

Images for global brain morphology (Figure 2B, Figure 3B, Figure 5A, and Figure 6D) were obtained using a Zeiss Axiozoom V16 macroscope, equipped with a digital camera (AxioCam HRm) using a 16x (pixel size: 4.037 μm), 63x (pixel size: 1.032 μm), or 80x objectives (pixel size: 0.813 μm). Images for NG2, PDGFRa, or CC1 quantifications were acquired using a Zeiss spinning-disk confocal CSU-W1 microscope with 25x oil objective (Z-plane step size: 0.68 μm). The first 15 z-planes were projected and shown in the figures.

Thickness of the corpus callosum (labelled with MBP or NF) was measured manually using “Measure” function in ImageJ/Fiji^46^. For MBP or NF labelling in the cortex, the first 15 z-planes of each image were projected using “Z Project” function. An ROI was drawn around the region of interest. Background was subtracted using “Subtract Background”. The raw integrated density of the signal was measured using “Measure” function. The value was divided by the area of the ROI in µm² and reported. All the steps were performed in Fiji.

The maximum intensity of MBP labeling was measured manually. The first 15 z-planes of each image were projected using “Z Project” function. A line passing crossing the center of corpus callosum vertically was drawn. Background was subtracted using “Subtract Background”. The maximum intensity of the signal was measured using “Measure” function in ImageJ/Fiji.

A home-made plugin was used to quantify the number of *Sox10*-GFP, NG2, PDGFRa, and CC1 cells. Since NG2 is a chondroitin sulfate proteoglycan which is expressed on the cell surface of OPCs^47^, it is difficult to segment the NG2 labeling and count them automatically. Therefore, in this plugin, Hoechst and GFP signals are segmented using the StarDist plugin in Fiji. The volume of the signal is filtered by the minimum and maximum values selected by the user. For GFP signal, there is a threshold for minimum intensity. This minimum intensity is measured manually by the user based on the comparison of the signal of the real GFP versus the background. The detected nuclei colocalized with segmented GFP signals are considered as the OL lineage cells (*Sox10*-GFP cells). Then, the plugin dilates the area of each segmented nucleus by 1 μm and measures “meanIntCor” for NG2, PDGFRa, or CC1, based on the following formula:

*meanlntCor = meanlnt - bg*

“meanInt” is the mean intensity of the signal of interest (NG2, PDGFRa, CC1) for dilated nucleus. “bg” is the mean intensity of the minimum signal in the projection.

Then, a threshold is selected for “meanIntCor” value to count the number of OPCs (NG2 or PDGFRa) or OLs (CC1).

### Single cell transcriptomic data analysis

Metadata and expression matrix from sequencing of single cells of the oligolineage at E13.5, P7 and juvenile/adult mouse (NCBI GEO Series: GSE95194; Marques *et al.* (2018)^20^) were downloaded as .RDS from “https://cells.ucsc.edu/?ds=oligo-lineage-dev”. R version 4.1.0 (2021-05-18) and Seurat_4.1.0 packages were used for analysis. Data were normalized and scaled before running the principal component analysis (PCA). Cell cycle scoring did not show any bias that could impact the PCA analysis. Further analysis (Figure S2) showed the batch effect between the developmental and juvenile/adult data that were not generated at the same time (Marques *et al*, 2016^28^). In order to remove the resulting technical variations, we used Harmony (v 0.1.1) to integrate the data. The results matrix was used as input for Uniform Manifold Approximation and Projection (UMAP) and the identification of cell clusters thanks to the shared nearest neighbor (SNN) graph (FindNeighbors and FindClusters R functions). In order to determine the useful resolution, we used clustree (v_0.5.0) and chose clusters generated at the resolution of 0.8. Overrepresentation and Gene Set Enrichment analyses were performed using clusterProfiler (v4.0.5). Assignment of clusters to cell types was achieved using the database msigdbr (R package v_7.5.1, cell type signature gene sets (M8) from *mus musculus*) and the data from the original Marques *et al.* (2018)^20^ analysis. The Gene Ontology (GO) database was used to compare *C1ql1^pos^* versus *C1ql1^neg^* cells.

## Acknowledgements

We would like to thank Lamia Bouslama-Oueghlani and Noemie Ades from ICM, Paris for their scientific suggestions, Philippe Mailly and Heloise Monet from CIRB Imaging facility (ORION) for their contribution in developing the home-made plugins to quantify the smFISH and immunofluorescence data, William D. Richardson from UCL, London for *Sox10^DTA^*mice and scientific suggestion, and CIRB administration and animal facility personnels. This work was supported by funding from: European Research Council ERC consolidator grant SynID 724601 (to FS), INCA PEDIAHR21-014 (to FS). SM received a PhD thesis funding from la Ligue Nationale Contre le Cancer and Ecole des neurosciences de Paris il-de-France (ENP). Since 2019, ENP has changed to la Fondation des Neurosciences Paris (FNP).

---

**Figure S1.**
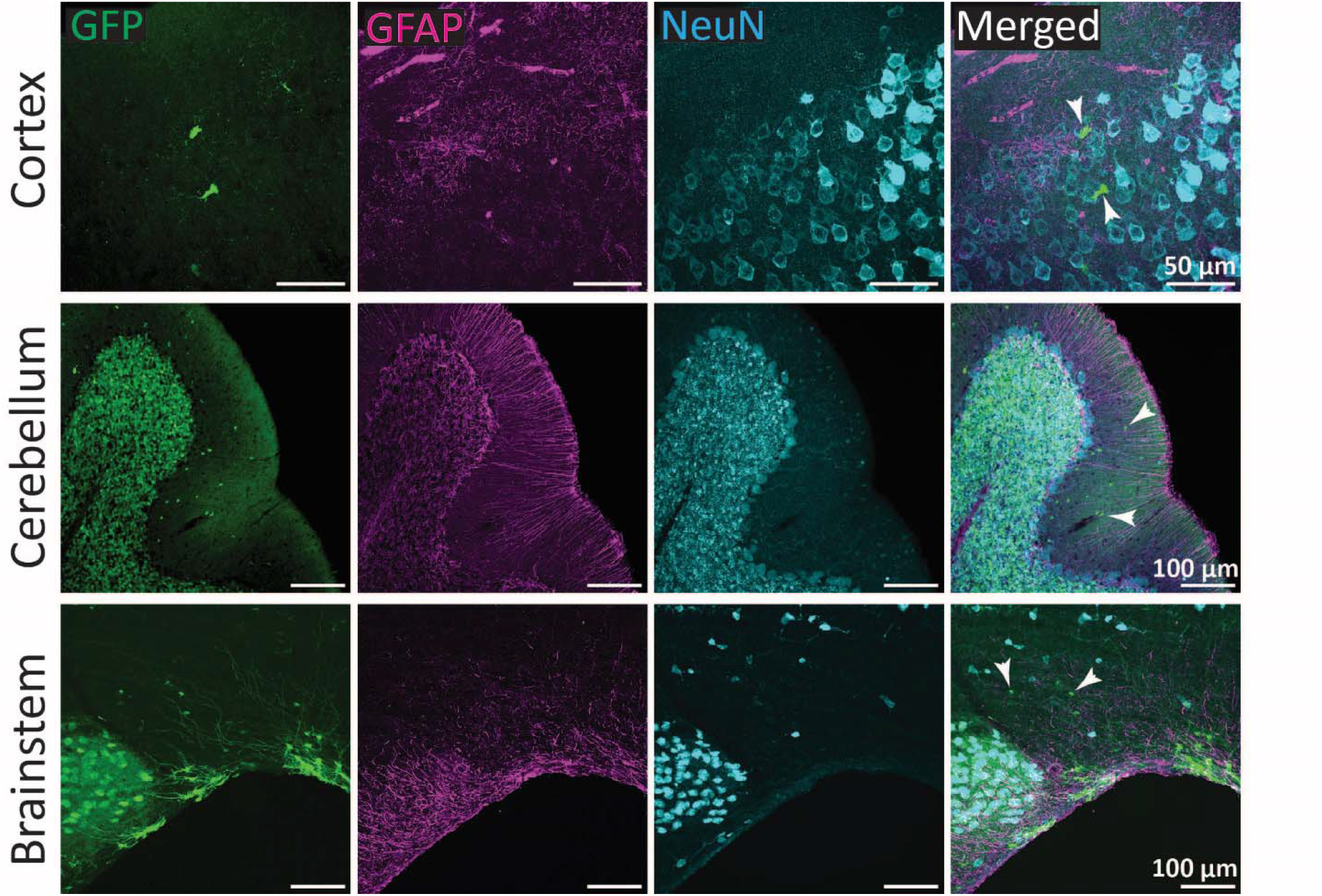
GFP-expressing cells in *C1ql1^Cre^; R26^Cas9-GFP^* brains are not astrocytes or neurons (except granule cells and inferior olivary neurons). GFP-expressing cells, astrocytes and neurons were visualized using co-immunolabeling for GFP (green), Glial fibrillary acidic protein (GFAP), and NeuN respectively, in sagittal brain sections from *C1ql1^Cre^; R26^Cas9-GFP^* animals at P30. The granule cells in the cerebellum and inferior olivary neurons in the brainstem are co-labeled with GFP and NeuN. GFP^pos^ GFAP^neg^ NeuN^neg^ cells (white arrowhead) are observed in all analyzed regions of the brain. Scale bars = 50 µm for cortex, Scale bars = 100 µm for cerebellum and brainstem.

**Figure S2.**
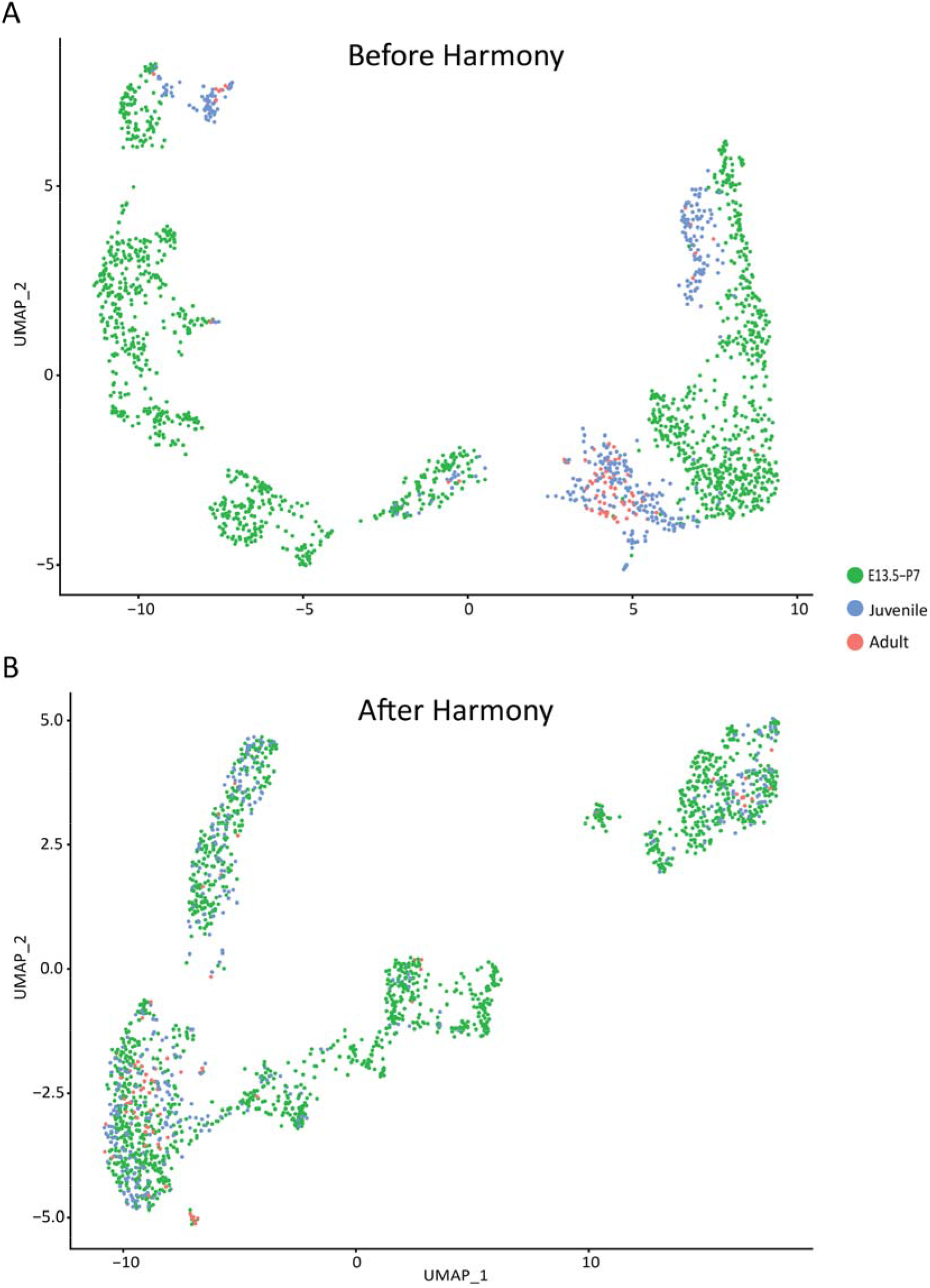
UMAP analysis of the single cell transcriptomic raw data from Marques *et al.*^20^: A) Data that were obtained in two separate studies (Juvenile and Adult Marques *et al.* 2016^28^ and E13.5 and P7 in Marques *et al.* 2018^20^) appeared initially as separate in the UMAP representation. B) After the integration of the data using Harmony, the batch effect was absent.

**Figure S3.**
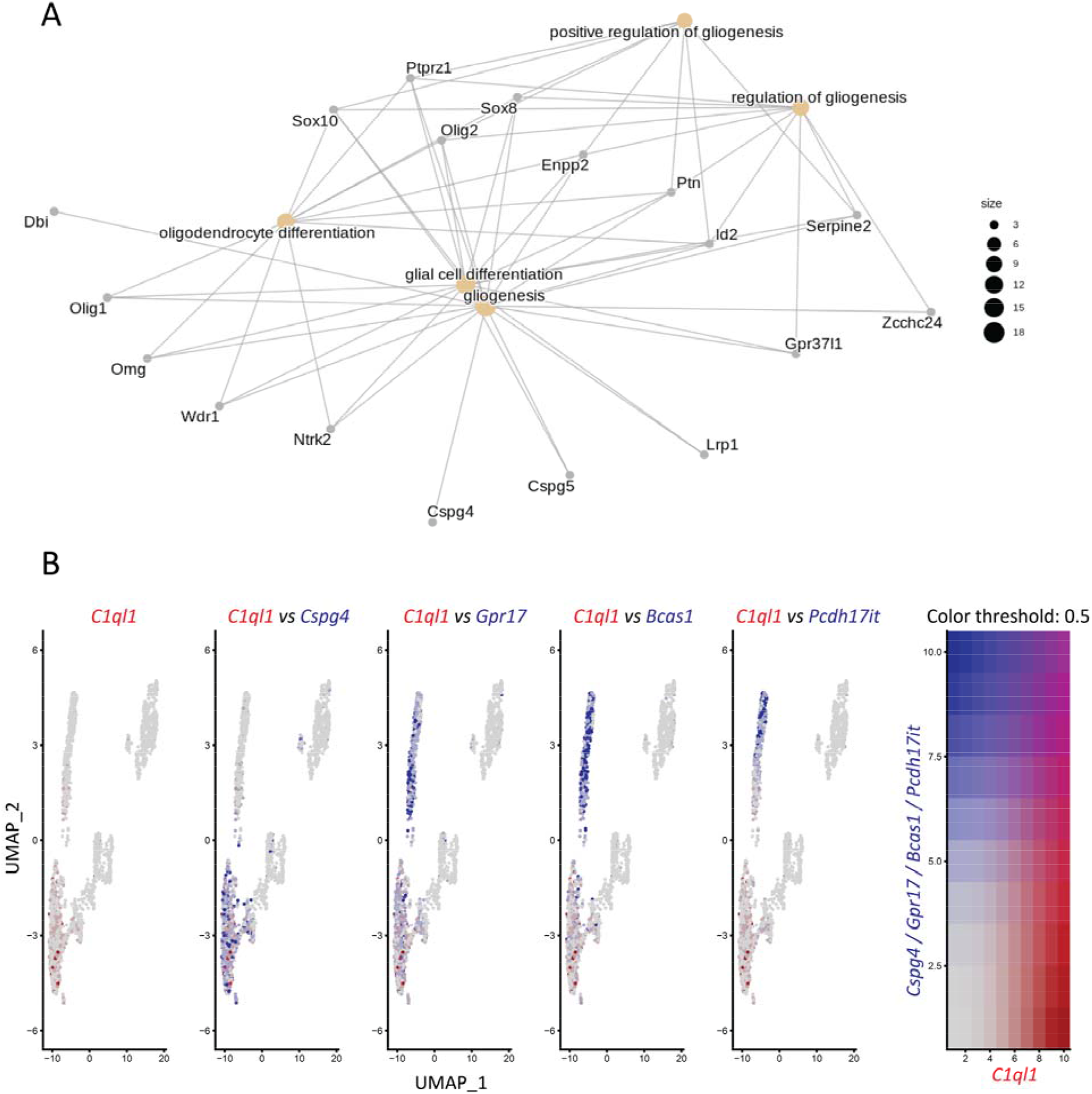
*C1ql1* marks a differentiation stage distinct from the COP and NFOL stages. A) Network analysis shows the genes co-enriched with *C1ql1* in E13.5, P7 and juvenile adult *Pdgfra*/GFP^pos^ cells (Reanalysis of raw data from Marques *et al.*^20^). B) New analysis of the single cell transcriptomic raw data from Marques *et al.*^20^ shows the overlap of the expression of *C1ql1* (*first left panel*) with the OPC marker *Cspg4* (coding NG2), but not with the COP and NFOL markers *Gpr17*, *Bcas1*, and *Pcdh17it*. UMAP: uniform manifold approximation and projection.

